# CellBinDB: A Large-Scale Multimodal Annotated Dataset for Cell Segmentation with Benchmarking of Universal Models

**DOI:** 10.1101/2024.11.20.619750

**Authors:** Can Shi, Jinghong Fan, Zhonghan Deng, Huanlin Liu, Qiang Kang, Yumei Li, Jing Guo, Jingwen Wang, Jinjiang Gong, Sha Liao, Ao Chen, Ying Zhang, Mei Li

## Abstract

In recent years, cell segmentation techniques have played a critical role in the analysis of biological images, especially for quantitative studies. Deep learning-based cell segmentation models have demonstrated remarkable performance in segmenting cell and nucleus boundaries, however, they are typically tailored to specific modalities or require manual tuning of hyperparameters, limiting their generalizability to unseen data. Comprehensive datasets that support both the training of universal models and the evaluation of various segmentation techniques are essential for overcoming these limitations and promoting the development of more versatile cell segmentation solutions. Here, we present CellBinDB, a large-scale multimodal annotated dataset established for these purposes. CellBinDB contains more than 1,000 annotated images, each labeled to identify the boundaries of cells or nuclei, including 4’,6-Diamidino-2-Phenylindole (DAPI), Single-stranded DNA (ssDNA), Hematoxylin and Eosin (H&E), and Multiplex Immunofluorescence (mIF) staining, covering over 30 normal and diseased tissue types from human and mouse samples. Based on CellBinDB, we benchmarked seven state-of-the-art and widely used cell segmentation technologies/methods, and further analyzed the effects of four cell morphology indicators and image gradient on the segmentation results.

## 1 Introduction

Cell/nuclear staining technology enhances the visualization of cell or subcellular resolution level in microscope images, thereby contributing to quantitative analysis in biomedical research[1]. As microscopy advances to study complex biological structures in great detail and produce rich, dense images[2], there is an urgent need for automated methods to extract cellular information by accurately segmenting cells and nuclei[1, 3], especially important in spatial transcriptomics[4]. Despite the significant progress made by deep learning methods in segmenting cell and nuclear boundaries in limited latent features of some specific images[5, 6, 7, 8, 9, 10, 11, 12, 13, 14, 15], challenges remain in developing universally applicable models. One of the key obstacles is the lack of large and diverse annotated image datasets that can support the training and evaluation of robust universal models[16, 17, 18, 19].

Previous works have made significant contributions by releasing several public datasets for training and evaluating deep learning models, although all important advances, they are limited in their ability to simultaneously meet the requirements for large-scale, multiple staining techniques, and diverse tissue types, thus unable to fulfill the needs for universal models. Table 1 shows the annotated stained datasets that have been actively used by the research community in recent years. As can be seen, some datasets are limited in scale or tissue type richness, such as MoNuSeg[16], IEEE_TMI_2019[20] and the fluorescence image dataset published by Kromp et al. in 2020[21], all of which contain fewer than 100 images. A slightly larger dataset Lizard[22] only consists of 1 tissue type. Table 1 also shows that most of the mentioned datasets are based on H&E staining, since H&E staining is the most common type of staining in routine pathology[23]. Followed by Immunofluorescence (IF) staining, which is more frequently employed in research settings. NuInsSeg[23] is limited to a single staining type of H&E. Similarly, BBBC039[24] is limited to Hoechst, a membrane-permeable fluorescent dye. Additionally, there are also three large-scale and more diverse datasets. The Data Science Bowl 2018 featured a dataset (BBBC038v1)[19] with 37,333 manually annotated cell nuclei, however, datasets commonly utilized for training and benchmarking segmentation models in stained cell images are still significantly larger by comparison. TissueNet[8] emphasizes immunostaining while lacking histological staining and nucleic acid staining. NeurIPS 2022 dataset[17] aims to achieve richness in image modalities, with constrained quantity in each image type, especially in the case of stained images.

Recently, many efforts have focused on developing universal methods and demonstrating robust performance on unseen datasets[6, 7, 8, 10, 26, 27]. SAM[26], a method based on the self-attention mechanism, has the capability of Zero-shot generalization of new image distributions and tasks. Cellpose1[6] is based on the U-Net architecture, which can precisely segment cells from diverse image types without model retraining or parameter adjustments. Cellpose3[7], an improved version, specializes in out-of-box segmentation of noisy, blurred or under-sampled images. DeepCell[8], a deep learning algorithm for accurate whole-cell segmentation that achieves human-level segmentation performance by combining a ResNet50 backbone network and a feature pyramid network. MEDIAR[27] emerged as the state of the art (SOTA) in the NeurIPS 2022 multimodal cell segmentation competition. StarDist[10] utilizes star-convex polygons to represent cell shapes, enabling accurate cell localization even under challenging conditions, especially when dealing with overlapping cells. However, the diversity of currently available models and the inconsistency of segmentation quality metrics make it difficult to evaluate their relative performance based on literature descriptions[28]. Therefore, it is necessary to evaluate these universal algorithms on a new unseen annotated dataset.

In this study, we present a new large-scale, multimodal annotated dataset, CellBinDB, containing images of four staining types (DAPI, H&E, ssDNA, and mIF) derived from over 30 human and mouse tissues. The primary statistic of CellBinDB is presented in the last row of Table 1. Unlike previous efforts that may be limited in tissue type coverage[21, 22], CellBinDB includes a wide range of both normal and diseased tissues, making it one of the most comprehensive tissue-type datasets available. The images included in the dataset were obtained from the 10x Genomics platform, as well as from new experiments based on Stereo-seq technology. Given that manual annotation is a time-intensive task, it is significantly limiting the scale of datasets. Besides, model annotation methods are affected by model style and cannot guarantee accuracy. To balance quality and efficiency in annotating this multimodal dataset, we used a combination of manual and semi-automatic annotation strategies. To eliminate potential biases in reference model[4], the model segmentation results were manually revised by a trained team of professional annotators and then checked by experts, ensuring that all annotations passed through two rounds of review. We make CellBinDB available to the research community and evaluate the performance of some general models on the dataset, providing recommendations for the selection of cell segmentation models in different scenarios.

**Table 1:** Publicly available annotated segmentation datasets.

| Dataset | Staining type | Image tiles | Cell | organs | Tile size(pixels) |
| --- | --- | --- | --- | --- | --- |
| MoNuSeg[16] | H&E | 44 | 28,846 | 9 | 1000 × 1000 |
| IEEE_TMI_2019[20] | H&E | 80 | 25,645 | 8 | 512 × 512 and 1000 × 1000 |
| Kromp et al.[21] | immuno \DAPI | 79 | 7,813 | 5 | 550 × 430 to 1360 × 1024 |
| Lizard[22] | H&E | 238 | 495,179 | 1 | 512 × 512 |
| NuInsSeg[23] | H&E | 665 | 30,698 | 31 | 512 × 512 |
| BBBC038v1[19] | DAPI\Hoechst\H&E | 841 | 37,333 | unknown | 256 × 256 |
| BBBC039[24] | Hoechst | 200 | 23,165 | unknown | 520 × 696 |
| TissueNet[8] | immuno | 3,200 | 1,300,000 | 9 | 512 × 512 |
| NeurIPS 2022 cell segmentation competition dataset<sup>1</sup>[25] | Jenner–Giemsa \immuno | 713 | 134,710 | unknown | 512 × 512 to 4096 × 4096 |
| **CellBinDB<sup>2</sup>** | **DAPI \ssDNA \H&E \mIF** | **1044** | **109,083** | **35** | **256 × 256 or 512 × 512** |

<sup>1</sup> This study focuses on cell segmentation of stained cell images, NeurIPS 2022 cell segmentation competition dataset only counted stained images.
<sup>2</sup> The last row of the table represents CellBinDB introduced in this work.

## 2 Results

### 2.1 Dataset

In this study, we propose CellBinDB, a dataset comprising 1,044 annotated microscopy images and 109,083 cell annotations. This dataset contains four staining types: DAPI, ssDNA, H&E, and mIF (Figure 1a). CellBinDB encompasses samples derived from human and mouse species, covering over 30 histologically diverse tissue types, including disease-relevant tissues (Figure 1b). The images in CellBinDB come from two sources: 844 mouse images were from in-house experiments based on Stereo-seq technology, and 200 human images obtained from the open-access platform 10x Genomics. We annotate all images in CellBinDB and offer two types of image annotations: semantic and instance masks (Figure 1c). Annotation is a combination of manual and semi-automatic methods, with 60% manual annotation and the rest from semi-automatic annotation (Figure 1f). The annotation workflow is shown in Figure 1g, and all annotations double-checked by experts to ensure quality. Additional details are in the Methods section. CellBinDB has diverse features that make it suitable for training generalized segmentation models. To visualize the image features of CellBinDB, we applied t-distributed stochastic neighbor embedding (t-SNE)[29, 30] to neural network-learned image features, revealing clusters that generally align by staining type (Figure 1d). The ssDNA and DAPI staining types show similar features, while there are significant differences between images of the same staining type from the two sources. CellBinDB exhibits a wider feature distribution, compared with previous datasets which share overlapping staining types. It encompasses features from most public datasets (Figure 1e). Supplementary Table 1 provides further details about CellBinDB.

**Figure 1.**
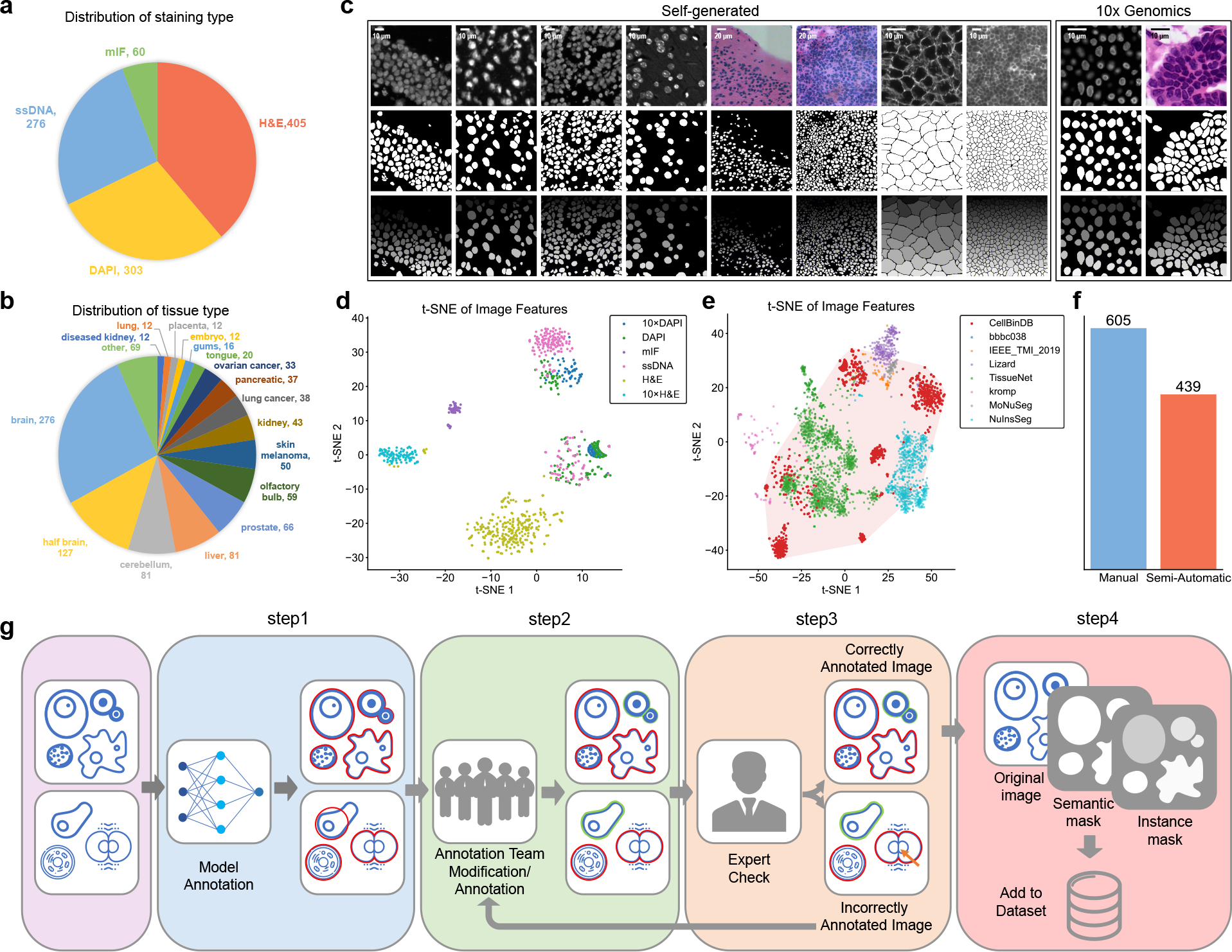
CellBinDB overview. a, Distribution of staining types in CellBinDB. b, Distribution of tissue types in CellBinDB, where tissue types with less than 10 samples are included in other, for details, see supplementary table 1. c, Examples of CellBinDB images and their two types of (semantic and instance) ground truth annotations, with 10 *µ*m scale bar from left to right columns 1, 2: ssDNA, columns 3, 4: DAPI, columns 5, 6: H&E, columns 7, 8: mIF, column 9: 10x Genomics DAPI, column 10: 10x Genomics H&E. The first row is the original microscope image, followed by the semantic annotation mask and instance annotation mask. d, Scatter plot of t-SNE demonstrates the diverse spread of data by different staining types and sources. e, Scatter plot of t-SNE demonstrates the diversity of CellBinDB compared to previous datasets. f, The number of manual and semi-automatic annotations in CellBinDB. g, The dataset annotation process includes four steps: 1.model annotation, 2.annotation team modification/re-annotation(depends on the model annotation results), 3.expert review, go to the next step if the annotations are correct, otherwise return to the second step for modification, 4. add the original image and the two masks to the dataset.

### 2.2 Benchmark performance

We evaluated several widely recognized segmentation models on CellBinDB, including specialized cell segmentation models: Cellpose1[6], Cellpose3[7], StarDist[10], DeepCell[8], MEDIAR[27, 25], a model for general segmentation: SAM[26], and a software widely used in biomedicine and life sciences: CellProfiler[9]. Among them, Cellpose1, Cellpose3, StarDist, and DeepCell are models based on U-Net, MEDIAR and SAM are models based on Transformer, and CellProfiler is a model based on machine learning. Since the models were trained on different training sets, a fair evaluation was conducted using CellBinDB, a dataset that none of the models had been trained on. The evaluation on CellBinDB was first performed on the entire dataset, followed by separate evaluations on each staining type.

#### 2.2.1 Evaluation results on the entire dataset

The initial objective was to assess the capacity of each model to perform segmentation on multimodal cell images (Figure 2a). The experimental results demonstrate that most of them exhibited excellent performance except for CellProfiler and DeepCell. Of the models evaluated, Cellpose3 demonstrated the most optimal performance and was the most highly recommended (precision: 0.82, recall: 0.61, F1 score: 0.70, dice: 0.72). In contrast, the performance of Cellpose1, with a similar architectural design, was less impressive in terms of a lower F1 score than Cellpose3. It is hypothesized that this inferior performance is attributable to the smaller and less diverse training dataset employed for Cellpose1 in comparison to Cellpose3. Similarly, DeepCell demonstrated suboptimal overall performance due to the limited diversity and lack of variability in its training dataset, which consisted exclusively of fluorescent staining images. Furthermore, the machine learning-based CellProfiler model demonstrated the poorest performance among the evaluated models.

**Figure 2.**
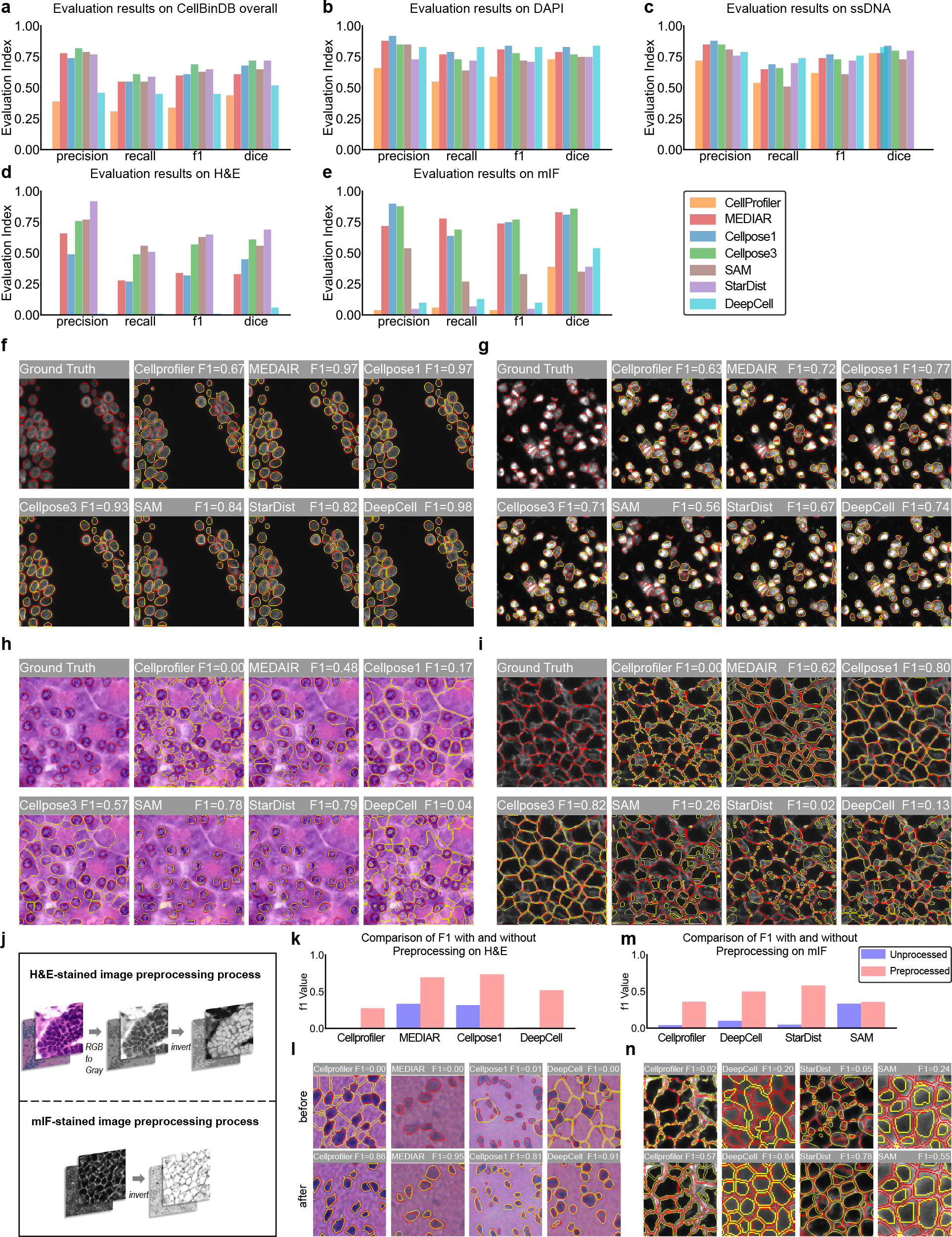
Evaluation of model performance on the entire dataset and by staining type. **a**, Evaluate model performance on the entire dataset, with bar charts of precision, recall, F1 score, and dice. **b**, Model performance evaluation results on DAPI-stained images. **c**, Model performance evaluation results on ssDNA-stained images. **d**, Model performance evaluation results on H&E-stained images. **e**, Model performance evaluation results on mIF-stained images. **f-i**, Examples of segmentation results of seven cell segmentation models on four stainings (DAPI, ssDNA, H&E, mIF). **j**, H&E and mIF-stained image preprocessing process. H&E-stained images include two steps: grayscale conversion and color inversion, while mIF-stained images only need to perform color inversion. **k**, Comparison of F1 scores before and after adding the preprocessing process for the model with improved performance of H&E-stained images. **l**, The segmentation results of the four models with the worst segmentation results on the H&E-stained image before and after image preprocessing. **m**, Comparison of F1 scores before and after adding the preprocessing process for the model with improved performance of mIF-stained images. **n**, The segmentation results of the four models with the worst segmentation results on the mIF-stained image before and after image preprocessing.

#### 2.2.2 Evaluation on four staining types respectively

We next evaluate the segmentation capabilities of each model on four staining types respectively. Figure 1 illustrates the diverse characteristics of images in CellBinDB. The evaluation of the models on the four staining types highlights the varying levels of difficulty across the stains and the strengths of each model. In detail, most models scored higher on DAPI (Figure 2b,f) and ssDNA (Figure 2c,g) stains. Notably, Cellpose1, DeepCell, and MEDIAR performed exceptionally well on these two stain types and are highly recommended. In contrast, significant variability was observed in model performance on H&E (Figure 2d,h) and mIF (Figure 2e,i) stains. On H&E-stained images, StarDist, SAM, and Cellpose3 performed well, whereas other models showed poor results or even failed. We attribute StarDist’s superior performance to its specialized weights trained for H&E-stained images, while SAM and Cellpose3 benefited from their large training datasets, enabling them to effectively segment H&E images. The poor performance of CellProfiler and DeepCell likely stems from their incorrect interpretation of the foreground and background (Figure 2h). mIF is a membrane stain that has the opposite color characteristics compared to DAPI and ssDNA, showing that the inside of the cell is black. On mIF-stained images, Cellpose1, Cellpose3, and MEDIAR are the most recommended. While SAM was able to process mIF images, its performance was relatively poor. The remaining models failed to effectively perform cell segmentation on mIF images because they mistakenly segmented the bright cell membranes as cells. Overall, H&E-stained and mIF-stained images pose greater segmentation challenges than other stain types.

Considering the specificity of H&E-stained image, some models perform additional preprocessing of the image, such as converting RGB to grayscale, or processing color deconvolution[31], to make the image features more consistent with the fluorescent stained image. We applied these preprocessing steps and re-evaluated the bottom four models when evaluated without preprocessing, finding that all four models demonstrated significant performance improvements (Figure 2k,l). DeepCell benefited the most; without preprocessing, it could barely segment H&E images. Cellprofiler, MEDIAR, and Cellpose1 also improved to varying degrees. This suggests that models specifically designed for fluorescent images can be extended more effectively to RGB image segmentation through preprocessing (RGB to grayscale conversion and then color inversion). After the above preprocessing, we recommend using Cellpose1, Cellpose3 or MEDIAR for cell segmentation on H&E-stained images. Similarly, this approach can be extended to mIF staining, with several previously underperforming models benefiting, particularly StarDist (Figure 2m,n), but still did not outperform Cellpose1, Cellpose3, and MEDIAR. The preprocessing process is shown in Figure 2j. Moreover, SAM demonstrates the capacity to process a range of stain types with consistent efficacy, although its performance is not as robust as that of models tailored for cell segmentation. This is believed to be due to the training data used for SAM, which is likely to contain a significant number of images that are not cells. In the absence of predefined settings or preprocessing, it is recommended to utilise SAM as a model capable of multimodal cell image segmentation.

### 2.3 Factors affecting cell segmentation performance

Next, we analyzed the factors that affect the model’s ability to segment cells from both biological and non-biological perspectives.

#### 2.3.1 Evaluation of impact of cell morphology

We evaluated a series of cell morphology metrics, including the cell morphology evaluation metrics in BIDCell[32] as well as an original metric, cell average distance, for evaluating cell density. The four metrics that exhibited the strongest correlation with the F1 score in the case of CellBinDB were found to be cell area, average distance, cell circularity and cell compactness. The results of the experiment indicate that an increase in the metrics is associated with an improvement in the performance of the segmentation. To illustrate, Cellpose1 demonstrated the most favourable performance in fluorescence staining (Figure 3a,c), whereas StarDist exhibited the most optimal results in H&E-stained images (Figure 3b,d). The full set of conclusions, which includes all models, can be found in the Supplementary Figure 1. The fluorescence staining images do not exhibit a notable distinction between the low and medium categories with respect to the indicator cellArea. This observation may be attributed to the intrinsic characteristics of the data, the cell areas of majority of images displaying cell areas concentrated within the region with lower values.

**Figure 3.**
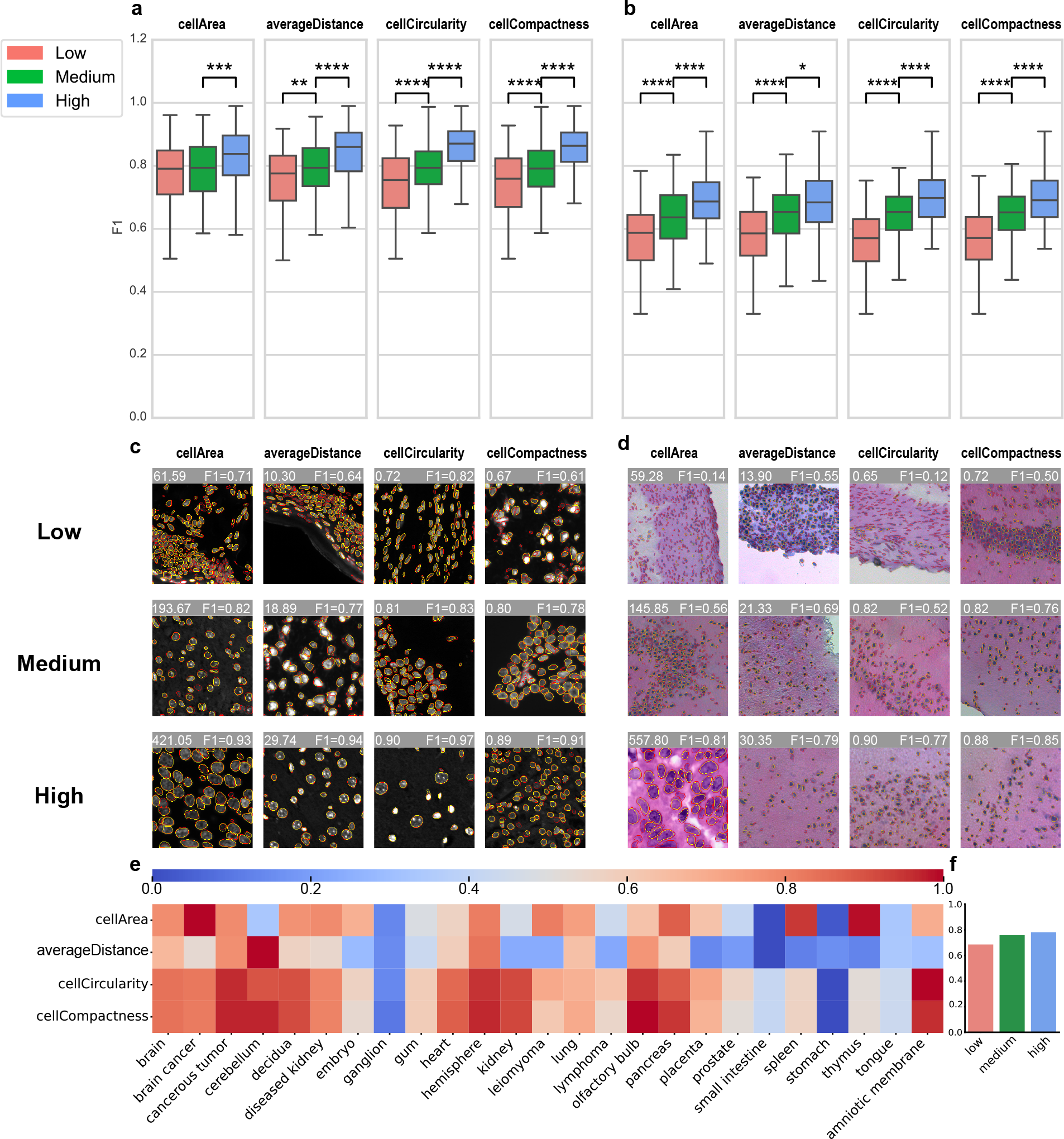
Evaluation of the impact of cell morphology. a, For fluorescent stained images (DAPI and ssDNA, Cellpose1 as an example), four indicators (cellArea, averageDistance, cellCircularity, cellCompactness) are used to evaluate the impact of cell morphology on segmentation performance. The vertical axis of the box plot is the F1 score, and the horizontal axis is the sample divided into three parts: low, medium and high according to the tertile of each indicator. And the significance mark line and p-value are added to the box plot. The number of “*” from 1 to 4 represents p-value less than 0.05, 0.01, 0.001 and 0.0001. b, Same as subfigure a, results on H&E-stained images. c, Fluorescent staining images display of low, medium and high instances under four indicators. d, H&E-stained images display of low, medium and high instances under four indicators. e, Display of normalized scores for the above four metrics by tissue type. f, According to the mean of the four cell morphology evaluation indicators of each image, it is divided into three groups: low, medium and high. The bar graph shows the relationship with the F1 score.

Cell morphology is directly related to tissue type. We classify the images in the dataset according to tissue type, calculate the above four indicators, and then normalize them. These results are presented in Figure 3e, which provides an overview of the difficulty level of cell segmentation for different tissue types and indicates the potential challenges encountered in cell segmentation for each tissue type based on specific cellular features. Finally, we computed the mean value of the aforementioned four normalized cell morphology indicators. Based on these mean values, we stratified the image samples into low, medium, and high groups and further explored the correlation between the mean values of normalized cell morphology indicators and the F1 score. The results indicated a positive correlation between mean values and F1 scores (Figure 3f). This finding suggests that the four selected indicators may, to some extent, reflect the difficulty of cell segmentation tasks.

#### 2.3.2 Evaluation of Image Quality Factors on Model Segmentation Performance

Morphological metrics are used to assess the impact of biological factors on samples, this experiment investigated the influence of image quality factors. Many deep learning models incorporate gradient information into their loss functions, such as the flow field in Cellpose1. The objective of this experiment is to investigate the impact of cell image gradients on model segmentation performance. The results demonstrate that, with regard to CellBinDB, the gradient in cell images does, in fact, exert an influence on the outcomes of the segmentation process.

The relationship between the image gradient magnitude calculated by the Sobel operator and the F1 score shows that most algorithms perform better on cell images of high-gradient magnitude, while their performance on low-gradient magnitude images deteriorates significantly (Figure 4a,b,c). This suggests that high-gradient magnitude images often provide more edge information, facilitating cell segmentation models to perform better in accurately locating cell boundaries. Meanwhile, we also observed that the performance difference between high-gradient and medium-gradient images is relatively small. This suggests that gradient variation may not be the sole determining factor in segmentation performance, with other factors such as noise and the complexity of cell morphology potentially playing a significant role. It can therefore be concluded that higher gradient magnitude is beneficial to cell segmentation models, as they assist in edge detection and feature extraction.

**Figure 4.**
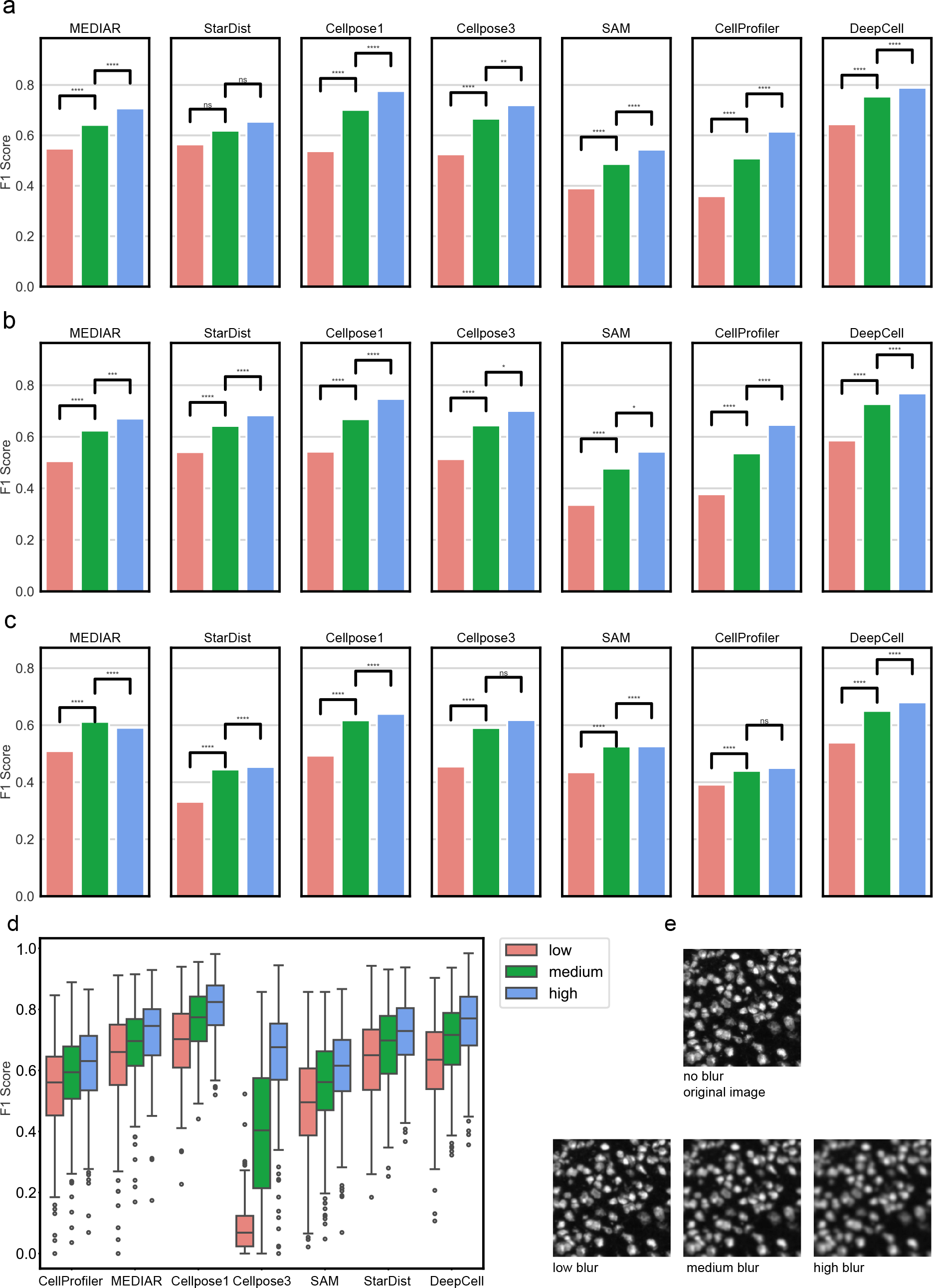
F1 Score vs Cell Gradient Groups for Different Algorithms. **a**, DAPI-stained images, relationship between cell gradient and F1 score. The vertical axis of the box plot is the F1 score, and the horizontal axis is the sample divided into three parts: low, medium and high according to the tertile of cell gradient. And the significance mark line and p-value are added to the box plot. The number of “*” from 1 to 4 represents p-value less than 0.05, 0.01, 0.001 and 0.0001. **b**, Same as subfigure a, results on ssDNA-stained images. **c**, Same as subfigure a, results on H&E-stained images. **d**, Simulation Experiment Results. **e**, examples of low, medium and high Gaussian blurred images.

To validate the above conclusions, we conducted simulation experiments and applied varying degrees of Gaussian blur to ssDNA images to reduce the gradient magnitude (which are detailed in the Methods section). We generated cell images with different gradient characteristics using this method (Figure 4e), and grouped them into low, medium, and high gradient categories. Next, we applied all algorithms to segment the cells in these images and calculated the F1 scores for each algorithm across the different gradient groups. As shown in Figure 4d, the results indicated that for all algorithms, the low, medium, and high gradient groups corresponded to low, medium, and high F1 scores, respectively. This means that high-gradient cell images, with clearer edge details, generally yielded better segmentation outcomes, while low-gradient images were more difficult to segment, leading to lower F1 scores. These experimental results further validate our previous conclusions, namely that high-gradient cell images are easier to segment. By reducing gradients of the images, simulation of varying degrees of edge information loss observed a reduction of segmentation accuracy in low-gradient images. This demonstrates the critical role of gradients in cell segmentation.

## 3 Discussion

In this paper, we introduce CellBinDB, a comprehensive and large-scale multimodal annotated dataset designed to enrich the current cell segmentation data ecosystem and advance the development of universal cell segmentation models.

Based on CellBinDB, we evaluated the performance of cell segmentation models on a series of benchmark tasks. CellBinDB is entirely new and unseen for all the evaluated models. We first evaluated the robustness and generalization of the cell segmentation models on the entire dataset. Our results show that without retraining and fine-tuning, the non-deep learning method CellProfiler has limited generalization ability and its performance significantly lags behind deep learning models. In stark contrast, deep learning models, especially Cellpose3, show excellent segmentation and generalization performance, highlighting their superiority in this field. Considering the significant difference in different staining types and image features learned by various models from training set, we then evaluated the cell segmentation models on image subsets of four staining types separately. In this evaluation, we found the challenges of cell segmentation on H&E-stained images and mIF-stained images and subsequently proposed a corresponding solution. Based on the above cell segmentation tasks, we conducted a comprehensive evaluation of the model performance and provided model selection suggestions for different cell segmentation scenarios.

Subsequently, we explore the factors that affect cell segmentation results. First, from the biological factors of the samples, differences in cell morphology such as cell area, cell density, and roundness directly affect the difficulty of cell segmentation. We used four indicators that were positively correlated with F1 score to quantify this effect. Different tissue types are directly related to cell morphology. We provide the normalized scores of cell morphological indicators for each tissue in CellBinDB to quantify the difficulty of cell segmentation in different tissues. Through this evaluation, we can estimate the performance of the model by the tissue type of the segmented image. For example, we can predict that cell segmentation in tissues such as the small intestine, ganglion, and tongue will be challenging. Finally, the evaluation of the impact on image quality shows that high image gradient magnitude facilities cell segmentation. The error factors such as image noise, blur, etc. will affect image gradient, which is possibly due to the limitations of the optical system itself, the sample preparation process, and the uncertainty of imaging conditions. Therefore, laboratory personnel should strictly perform experimental operations to provide higher quality image data for downstream systems.

While this work has made a contribution, it is important to acknowledge its limitations. The dataset utilized in this study is confined to two-dimensional (2D) images despite the variety of images. In recent years, the popularity of three-dimensional (3D) images has increased, posing new and unique challenges for cell segmentation tasks. To address these emerging challenges, researchers have already developed algorithms capable of performing 3D cell segmentation. Acknowledging the importance of this advancement, we intend to delve into this direction and explore the potential of incorporating 3D images into CellBinDB in future research endeavors.

In conclusion, compared to any previous work, this work makes a contribution in diversity and scale of dataset, offering a comprehensive and extensive dataset in this work that serves as a valuable resource for advancing cell segmentation models. Nevertheless, this represents only a small step forward in the direction of universal model development. Given the complexities and nuances in the field, ongoing research and continuous expansion of datasets encompassing various staining types, tissue types, and imaging modalities are necessary. By continuing to build upon this preliminary progress, we can collectively work towards the ultimate goal of developing models that can generalize and adapt to diverse datasets, staining techniques, and tissue types in the field of cell segmentation.

## 4 Methods

The code library of this study is implemented in Python3[33], using numpy[34], scipy[35], pandas[36], skimage[37], opencv[38] and sklearn[30], and jupyter[39] was used to create a notebook version. The figures in this article were plotted using matplotlib[40] and seaborn[41], and formatted using Adobe Illustrator.

### 4.1 Data collection and annotation

#### 4.1.1 Image acquisition

CellBinDB includes 30 mouse tissues and 5 human tissues. The 5 different human tissue samples are from the open platform 10x Genomics, with download links provided in the Data Availability section. For two IF-stained images (human skin melanoma and human prostate cancer), only the DAPI channel was selected. The mouse tissues in this study were collected from 6-week-old C57BL/6 both female and male laboratory mouses. All experimental protocols for generating dataset adhered to ethical regulations regarding animal research. Fresh frozen samples were prepared and sectioned according to STOmics Stereo-seq Transcriptomics Set User Manual. Tissue slices of 10 *µ*m were attached to the chip surface and stained after the tissue fixation. During the imaging stage, the epi-bright field (color camera) mode was selected for H&E-stained tissues while the epi-fluorescence mode was selected for fluorescent-stained tissues. After completing the photography according to the manual requirements, the imageQC module of the StereoMap software was used for image quality control.

WSIs are generated by 1. a STOmics Microscope Go Optical equipped with Scanner Version 1.2.2, using 10×/0.75 NA and 20X/0.5 NA objective and Go Optical Scanner Ximea Mc124 for grayscale images and Go Optical Scanner Ximea Mc050 for RGB images. 2. a Motic PA53 FS6 Microscope equipped with PA53Scanner 1.0.0.14, using 10×/0.75 NA objective and PA53 FS6 SCAN S5LITE MONO for grayscale images and PA53 FS6 SCAN S5LITE for RGB images. We obtained whole slide images (WSIs) from the laboratory or open platforms and then cropped the size to 512×512-pixel or 256×256-pixel sizes. A biologist selected the most representative fields of view (FOV) for each WSI, ensuring that the selected FOV images were clear and suitable for creating ground truth.

#### 4.1.2 Image annotation

We trained a model on CellBinDB and trained a team of professional annotators to semi-automatically annotate the dataset proposed in this study through three steps: model segmentation, manual modification or re-annotation, and expert verification. The model we used for annotation was from Li, M. et al.[4] The model architecture was psaUnet contained 5 encoder blocks and 5 decoders blocks. In the preprocessing stage, median filtering was used to smooth the possible noise in the input image. And in the post-processing stage, operations such as corrosion, dilation, and watershed were performed. The preliminary results of the model segmentation were saved in Tagged Image File Format (TIFF) and then converted to JavaScript Object Notation (json) format and imported into Qupath 0.4.3 software[42]. In Qupath 0.4.3, the nuclei annotated by the model were visualized as polygons overlaid on the original nucleus images. If the model segmentation results were satisfactory, the annotation team obtained more accurate annotations by refining polygon outlines, deleting or adding polygons to modify the results of the model segmentation. Conversely, the annotation team re-annotated manually. The model we trained did not support cell membrane segmentation, therefore mIF-stained images must be annotated manually. Meanwhile the model annotation step was skipped, and the annotation team directly used Qupath 0.4.3 software to outline the cells with the polygon tool. These annotations were subsequently checked by experts. The incorrectly annotated images and corresponding modification suggestions were returned to the annotation team. Once the annotations were approved, the two types of masks: instance mask and semantic mask, were exported and converted into TIFF. Finally, the microscope images were added to the dataset along with the two forms of masks. Supplementary Table 2 provides detailed information such as the file name and staining type, tissue type, size, and data source for each image.

The annotation of cell is challenging due to out-of-focus or nuclei presenting with modified morphology during the slide preparation procedure. We defined the following criteria to annotate images:

- Naked eye is the highest standard and all cells that can be identified by naked eye are annotated ensuring no false negatives (FN) or false positives (FP).
- The entire cell must be completely covered without any omissions or obvious gaps.
- The annotated cell boundaries should be smooth without excessive jagged edges.
- Overlapping cells should be annotated separately.
- The segmentation should not be excessively fragmented. For elongated cells, do not split them into multiple individuals.

### 4.2 Model architecture and default parameters

SAM[26] claims that it has capability on Zero-shot generalization of new image distributions and tasks. The model_type of SAM utilizes ‘vit_b’ and applies SamAutomaticMaskGenerator to automatically generate masks without the need for external prompts.

Cellpose1[6] can segment multiple types of cells without requiring parameter adjustments, new training data or further model retraining. Cellpose1 uses the ‘cyto’ model, with the channels for grayscale images set to [0,0] and [1,3] forH&E images. The diameter is set to None.

Cellpose3[7] introduced a novel method for biological image that achieves efficient segmentation without the requirement for clean images. Cellpose3 uses the ‘cyto3’ model, with the channels for grayscale images set to [0,0] and [1,3] for H&E images. The diameter is set to None, and the ‘denoise_cyto3’ model is used for noisy images.

DeepCell[8] achieves human-level accuracy across a variety of tissues and imaging modalities while requiring no manual parameter tuning for the end user. DeepCell uses the ‘Mesmer’ model, with image_map set to 0.5 and compartment set to ‘nuclear’.

MEDIAR[27] stood out in the NeurIPS 2022 cell segmentation competition15 and achieved state of the art (SOTA). MEDIAR uses the provided from_phase2.pth model, with model_args configured as follows: ‘classes’ is set to 3, ‘decoder_channels’ is set to [1024, 512, 256, 128, 64], ‘decoder_pab_channels’ is set to 256, ‘encoder_name’ is set to ‘mit_b5’, and ‘in_channels’ is set to 3. The algo_params has ‘use_tta’ set to True.

StarDist[10] uses star-convex polygons to represent cell shapes, allowing accurate cell localization even under challenging conditions. StarDist uses the ‘2D_versatile_he’ model for HE images and the ‘2D_demo’ model for non-HE images.

Cellprofiler[9] is one of the earliest high-throughput cell image analysis platforms. Cellprofiler segmentation pipeline for non-H&E images is as follows: firstly, use IdentifyPrimaryObjects to identify cell objects, setting “Discard objects touching the border of the image” to No. Then, apply OverlayOutlines to outline the cells. Next, use ExpandOrShrinkObject to erode the cell boundaries, separating closely adjacent cells. MaskImage is utilized to convert the image to a mask and finally, save the image with SaveImage. For H&E images, the segmentation pipeline is similar but it starts with an additional step: convert the image to grayscale and then invert the colors before proceeding with the same steps as for non-H&E images.

### 4.3 Benchmark pipeline

Different from some previous work[28, 43, 44, 45], we designed a series of benchmarks to test the performance of each model without retraining. The method maximized the diversity and breadth of CellBinDB to evaluate the generalizability and robustness of each model. Unless otherwise specified, default parameters will be used. Each step of the benchmark targets a different cell segmentation scenario:

(1) Model evaluation on the entire dataset. First, we test each model on the entire dataset including all staining types and tissue types and evaluate the overall performance results using the metrics precision, recall, F1 score, and Dice. The step aims to recommend the model with the optimum overall performance when the user is segmenting a multimodal dataset or user id unfamiliar with their own data’s attributes.
(2) Model evaluation on different staining types. In this step, the dataset is classified into four staining types and the performance of each model is tested individually on each single staining type. This step is intended to furnish users with model recommendations for cell segmentation reliant on specific stain types, as well as to facilitate a comparison of the relative challenges imposed by different staining types.
(3) Evaluating the impact of cell morphology on the performance of cell segmentation models. To quantify cell morphology, we measured a series of metrics for each cell in the dataset, which represent different cell characteristics. We then selected several metrics (cell area, average distance, cell circularity, and cell compactness) with the greatest impact on the F1 score to explore the relationship between cell morphology and model performance. Initially, the values of these four indicators were computed for each cell, and the mean value for each image was determined. Subsequently, the images were categorized into three groups—low, medium, and high—based on tertiles, to assess whether there are significant differences in the F1 scores across these groups. Given the similarity in features between DAPI and ssDNA-stained images, these two types were analyzed together. In contrast, H&E-stained images were analyzed separately, and mIF-stained images were excluded from this experiment due to their unique membrane staining characteristics.
(4) Exploring the effect of cellular image gradients on model segmentation performance. First, we matched individual cells with ground truth and predicted results. Then, we calculated the gradient magnitude of each cell using the Sobel operator in OpenCV, along with its corresponding F1 score. Subsequently, cells were categorized into three groups—low, medium, and high gradients—to assess whether there were significant differences in F1 scores across these groups.

### 4.4 Metrics

#### 4.4.1 Segmentation benchmark

To evaluate models’ performance on benchmark pipeline, an evaluation protocol was used which is calculated in several steps. First, the overlap between each prediction and its closest ground truth object is quantified using the *IoU* as follows:

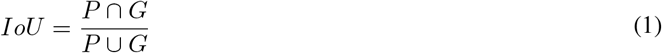

where *P* is prediction, *G* is ground truth. If the *IoU* between the prediction and the closest ground truth object is greater than 0.5, the ground truth object is considered to be correctly segmented. For all ground truth objects, the segmentation performance is then quantified using the *precision* and *recall* metrics given below:

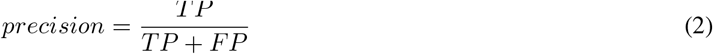

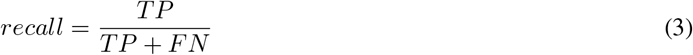

where *TP* is the number of true positives, *FP* is the number of false positives and *FN* is the number of false negatives. The *F1 score* is a metric used to assess the balance between *precision* and *recall*. It is the harmonic mean of *precision* and *recall*, providing a single score that encapsulate the characteristics of both parameters:

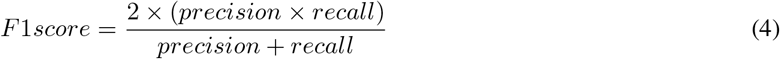

In addition, a pixel-level evaluation metrics are introduced:

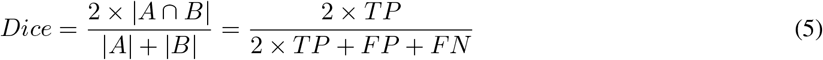

#### 4.4.2 Cytomorphological indicators

To explore the impact of cell morphology on segmentation performance, we measured a series of morphological metrics for each cell in the dataset and selected the ones with the greatest impact, including *cell area, average cell distance, cell circularity*, and *cell compactness*.

The *cell area* is the number of pixels contained in each cell. The *average cell distance* refers to the average distance between each cell and the center point of the nearest cell. *Cell circularity* is used to calculate the circularity of the cell shape. The closer the value is to 1, the closer the cell shape is to a circle. We used the cv2.convexHull() function in OpenCV to get the convex hull of each cell, used the cv2.arclength function to get the convex hull area, and then calculated the circularity according to the following formula:

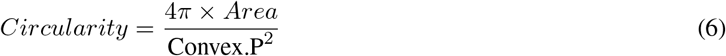

*Compactness* is the degree of compactness of the cell shape, calculated based on the cell area and cell perimeter. The closer the *compactness* is to 1, the more compact and circular the cell shape is; the closer the *compactness* is to 0, the more irregular and scattered the cell shape is:

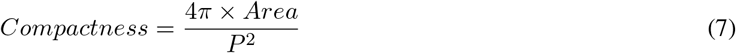

The difference between the above two indicators is that cell roundness uses the perimeter of the cell convex hull, while *compactness* uses the cell perimeter. If there are many irregular protrusions or depressions on the edge of the cell, the value of *compactness* will obviously deviate from 1. *Circularity* pays more attention to the roundness of the overall shape and is less affected by slight irregularities on the edge.

#### 4.4.3 Image gradient calculation

We used the ground truth mask to identify the cell regions in the cell images, then applied the Sobel operator from OpenCV (cv2) to compute the x and y gradients of the cell images, followed by calculating the gradient magnitude and average.

The formula for the gradient in the x-direction (*G*_*x*_):

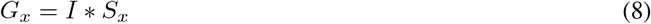

where *I* is the input image, and *S*_*x*_ is the Sobel kernel matrix for the x-direction:

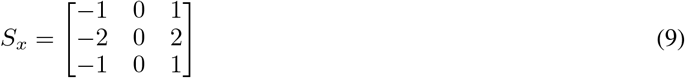

The formula for the gradient in the y-direction (*G*_*y*_):

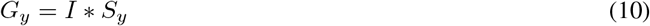

where *S*_*y*_ is the Sobel kernel matrix for the y-direction:

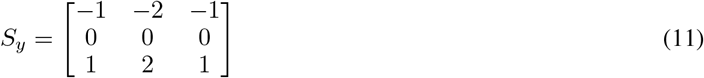

The gradient magnitude (*G*) is calculated as:

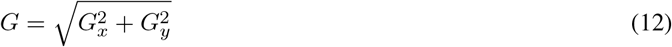

The average gradient magnitude is calculated as:

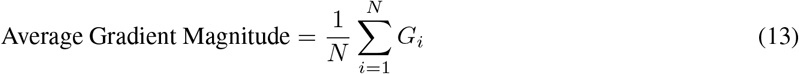

where *N* is the number of pixels in the image, and *G*_*i*_ is the gradient magnitude of each pixel.

### 4.5 Simulation experiments

We used several different Gaussian blur parameters and adjusted the gradients of these blurred images to create three categories: low, medium, and high gradients. Each category of images had varying levels of blur, aimed at simulating images with different gradient magnitudes in order to assess the impact of gradient on cell segmentation performance. The blurring was implemented using OpenCV’s GaussianBlur function. The parameters for low-intensity blurring were a (5,5) kernel size with sigmaX set to 1; for medium intensity, a (9,9) kernel size with sigmaX set to 2; and for high intensity, a (13,13) kernel size with sigmaX set to 3. After applying the blur, we calculated the average gradient magnitude of the images. The calculation formula for Gaussian blurring is as follows:

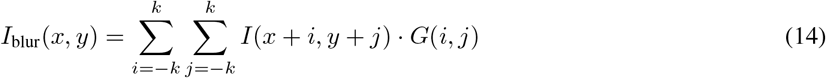

Where I(x+i,y+j) is the original pixel intensity at position (x+i,y+j), G(i,j)is the Gaussian kernel weight for the position (i,j) relative to the center.

## 8 Abbreviations

DAPI: 4’,6-Diamidino-2-Phenylindole
ssDNA: Single-stranded DNA
H&E: Hematoxylin and Eosin
mIF: Multiplex Immunofluorescence

## 5 Data availability

CellBinDB have been deposited into CNGB Sequence Archive (CNSA)[46] of China National GeneBank DataBase (CNGBdb)[47] with accession number CNP0006370, and also uploaded to zenodo[48].

The raw data of 5 human samples from 10×Genimics are available free of charge. The download links are as follows.

H&E human lung cancer: [FFPE Human Lung Cancer with Xenium Multimodal Cell Segmentation](https://www.10xgenomics.com/datasets/preview-data-ffpe-human-lung-cancer-with-xenium-multimodal-cell-segmentation-1-standard).

H&E human pancreas: [FFPE Human Pancreas with Xenium Multimodal Cell Segmentation](https://www.10xgenomics.com/datasets/ffpe-human-pancreas-with-xenium-multimodal-cell-segmentation-1-standard).

H&E human ovarian cancer: FFPE Human Ovarian Cancer Data with Human Immuno-Oncology Profiling Panel and Custom Add-on.

DAPI human skin melanoma: [Human Melanoma, IF Stained (FFPE)](https://www.10xgenomics.com/datasets/human-melanoma-if-stained-ffpe-2-standard).

DAPI human prostate Cancer: [Human Prostate Cancer, Adjacent Normal Section with IF Staining (FFPE)](https://www.10xgenomics.com/datasets/human-prostate-cancer-adjacent-normal-section-with-if-staining-ffpe-1-standard).

## 6 Code availability

Project name: CellBinDB: A Large-Scale Multimodal Annotated Dataset for Cell Segmentation with Benchmarking of Universal Models

Project home page: https://github.com/STOmics/cs-benchmark)

Operating system(s): Platform independent

Programming language: Python

Other requirements: Python 3.8 or higher License: MIT License

## 7 Ethics, consent and permissions

The human data in this study are public data collected from open platforms. The animal data produced by ourselves have passed the ethics review, and the ethics approval number is BGI-IRB A21001-T1.

## 9 Authors’ Contributions

Project administration and supervision: Mei Li, Ying Zhang

Algorithm development and implementation: Jinghong Fan, Huanlin Liu, Zhonghan Deng

Data provided: Yumei Li, Jing Guo

Data selection, processing and sorting: Can Shi

Data annotation: Can Shi, Jinjiang Gong, Jingwen Wang, Jinghong Fan, Ying Zhang, Yumei Li

Project coordination: Mei Li, Ying Zhang, Zhonghan Deng

Method comparisons: Can Shi, Jinghong Fan Code Testing: Can Shi, Jinghong Fan

Manuscript writing and figure generation: Can Shi, Jinghong Fan, Ying Zhang

Manuscript review: Can Shi, Jinghong Fan, Mei Li, Ying Zhang, Qiang Kang, Sha Liao, Ao Chen

## 11 Acknowledgments

We thank China National GeneBank for providing data storage support for this study.

