## Supplementary figures and images for "CellBinDB: A Large-Scale Multimodal Annotated Dataset for Cell Segmentation with Benchmarking of Universal Models"

### Supplementary Figure 1

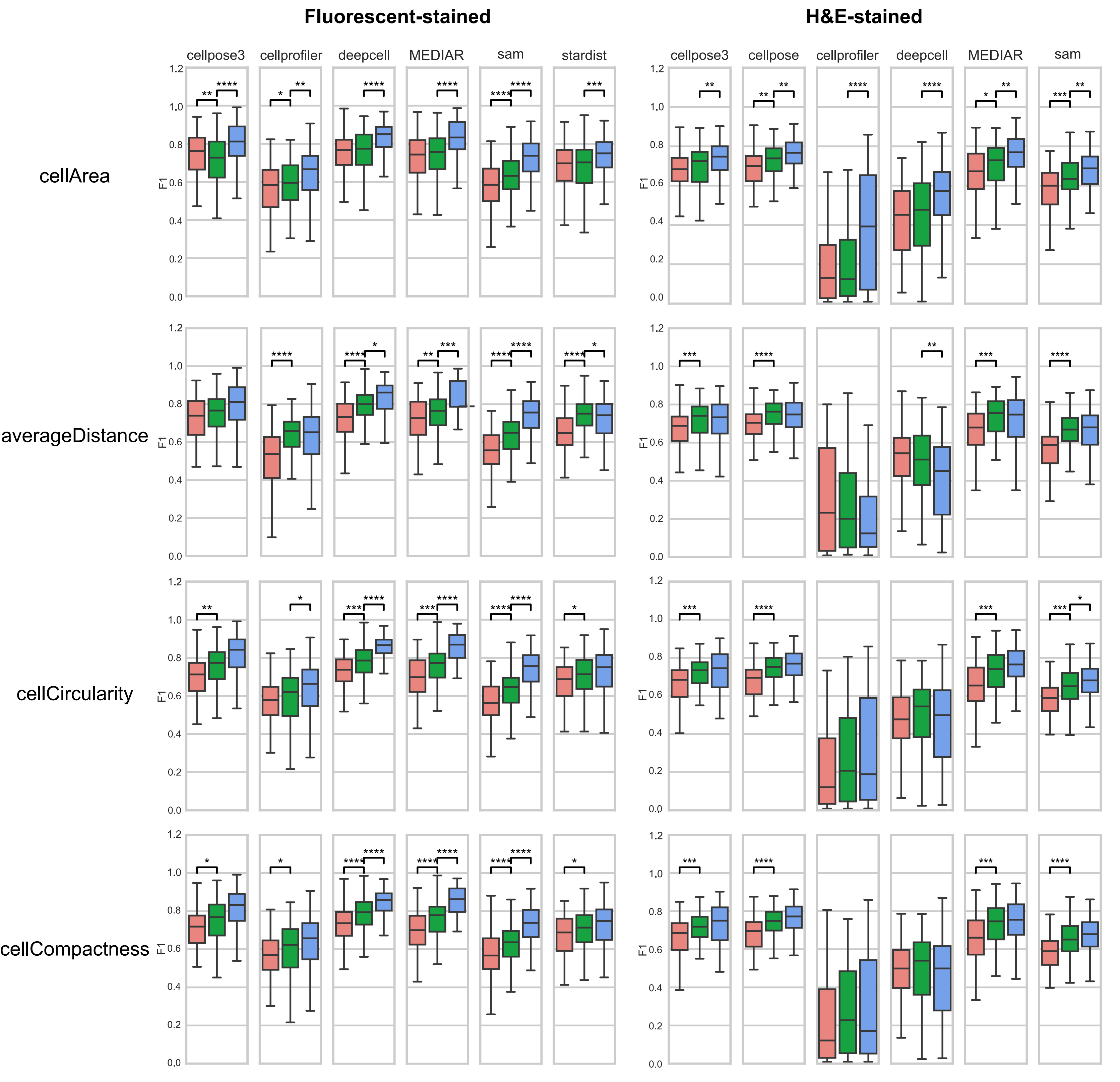
